# Optimizing oleaginous yeast cell factories for flavonoids and hydroxylated flavonoids biosynthesis

**DOI:** 10.1101/614099

**Authors:** Yongkun Lv, Mattheos Koffas, Jingwen Zhou, Peng Xu

**Affiliations:** Department of Chemical, Biochemical and Environmental Engineering, University of Maryland Baltimore County, Baltimore, MD 21250; National Engineering Laboratory for Cereal Fermentation Technology, Jiangnan University, 1800 Lihu Road, Wuxi, Jiangsu 214122, China; Department of Chemical and Biological Engineering, Rensselaer Polytechnic Institute, Troy, NY 12180; Jiangsu Provisional Research Center for Bioactive Product Processing Technology, Jiangnan University, 1800 Lihu Road, Wuxi, Jiangsu 214122, China

**Keywords:** Natural products, Flavonoids, Hydroxylation, Metabolic engineering, Oleaginous yeast

## Abstract

Plants possess myriads of secondary metabolites with a broad spectrum of health-promoting benefits. Up to date, plant extraction is still the primary route to produce high-value natural products, which inherently suffers from economics and scalability issues. Heterologous production in microbial host is considered as a feasible approach to overcoming these limitations. Flavonoid and its hydroxylated derivatives represent a diversified family of bioactive compounds, most prominently known as antioxidant and anti-aging agents. Oleaginous yeast is rich in hydrophobic lipid bodies and spatially-organized organelles, which provides the ideal environment for the regioselectivity and stereoselectivity of many plant-specific enzymes. In this report, we validated that *Y. lipolytica* is a superior platform for heterologous production of high-value flavonoids and hydroxylated flavonoids. By modular construction and characterization, we determined the rate-limiting steps for efficient flavonoids biosynthesis in *Y. lipolytica*. We evaluated various precursor pathways and unleashed the metabolic potential of *Y. lipolytica* to produce flavonoids, including the supply of acetyl-CoA, malonyl-CoA and chorismate. Coupled with the optimized chalcone synthase module and the hydroxylation module, our engineered strain produced 252.4 mg/L naringenin, 134.2 mg/L eriodictyol and 110.5 mg/L taxifolin from glucose. Collectively, these findings demonstrate our ability to harness oleaginous yeast as microbial workhorse to expand nature’s biosynthetic potential, enabling us to bridge the gap between drug discovery and natural product manufacturing.

## Introduction

Plant possess myriads of secondary metabolites with a broad spectrum of health-promoting benefits. Plant-derived natural products (PNPs) have been used to suppress tumor growth, inhibit retrovirus replication, treat metabolic disease and modulate cholesterol level in both animal and human tests ^1^. Up to date, plant extraction is still the primary route to produce PNPs ^1^. However, isolation of PNPs from their native sources is limited by low abundance and environmental, seasonal, and regional variations. Total chemical synthesis of complex PNPs often involves toxic catalyst and is commercially unsustainable due to low yield, safety concerns and strict GMP regulations ^2^. Genome-mining of plant metabolic pathways ^3–8^ and reconstruction of biosynthetic gene clusters (BGCs) in industrially-relevant microbes offer significant promise for discovery and scalable synthesis of plant natural products.

*E. coli* and *S. cerevisiae* have long been established as host strains to manufacture a large variety of plant natural products ^3, 7–14^. The recent development of oleaginous yeast platforms offers significant advantages over *E. coli* and *S. cerevisiae* (Table 1). The high precursor acetyl-CoA and malonyl-CoA flux along with the hydrophobic lipid bodies make oleaginous yeast a promising host to produce various natural products with complex structures. For example, *Y. lipolytica* is known to internalize substantial portion of carbon feedstock as lipids and fatty acids ^15, 16^, which provides the ideal amphiphilic environment for the catalytic function of many plant-derived enzymes. It has been recognized as a ‘generally regarded as safe’ (GRAS) organism for the production of organic acids, polyunsaturated fatty acids (PUFAs) ^17, 18^ and carotenoids ^19–22^ in the food and nutraceutical industry. Compared to *S. cerevisiae*, *Y. lipolytica* lacks Crabtree effects, which doesn’t require the co-feeding of ethanol for cell to grow. The low pH tolerance ^23^, strictly aerobic nature ^24, 25^ and versatile substrate-degradation profile ^25–27^ enable its robust growth from a wide range of renewable feedstocks. Genetic toolbox development has been expanding to protein expression ^28–30^, promoter characterization ^31–33^, YaliBrick-based cloning ^34, 35^, Golden gate cloning ^21, 36^, Piggyback transposon ^37^, genome-editing ^34, 38, 39^ and iterative gene integration ^40^, affording us a collection of facile genetic tools for streamlined and accelerated pathway engineering in oleaginous yeast species.

**Table 1.** Comparison of *E. coli*, *S. cerevisiae* and *Y. lipolytica* as chassis to produce plant natural products

|  |  |  |  |
| --- | --- | --- | --- |
| Expression platform | *E. coli* | *S. cerevisiae* | *Y. lipolytica* |
| Genetic tools | +++++ | +++++ | +++ |
| Genome annotation | +++++ | ++++ | ++++ |
| Acetyl-CoA/Malonyl-CoA/HMG-CoA flux | ++ | +++ | +++++ |
| P450 expression | + | +++ | +++++ |
| Substrate flexibility | ++++ | ++ | ++++ |
| Acid tolerance | +++ | ++ | +++++ |
| FDA safety | + | +++++ | +++++ |
| Hydrophobic lipid body environment | + | ++ | +++++ |

Flavonoids represent a diversified family of phenylpropanoid-derived plant secondary metabolites, with an estimated 10,000 unique structures ^41, 42^. They are widely found in fruits, vegetables and medicinal herbs and plants. Pharmaceutical studies and animal tests have demonstrated their anti-obesity, anti-cancer, anti-inflammatory, and anti-diabetic activities ^43, 44^. Flavonoids are among the phytochemicals with proven activity towards the prevention of aging-related diseases, including the treatment of nervous and cardiovascular diseases, Parkinson’s and Alzheimer’s disease etc. ^45^. These health-promoting benefits make flavonoids a distinct family of molecules to fight aging in personal care, nutraceutical industry and clinical trials. Considering the worldwide population with age older than 65 will triple in next 30 years (data from WHO, world health organization), there will be increasing interests and sustainable market demand globally in the near future.

To prepare the technological ground and enable microbial synthesis, various flavonoid pathways have been reconstituted in various microbial species including *E. coli* and *S. cerevisiae* recently, and the recombinant production of an array of molecules such as naringenin ^46, 47^, eriodyctiol ^48, 49^, resveratrol ^50–54^, pinocembrin ^49, 55^, anthocyanins ^56–58^, quercetin, kaempferol ^59^, silybin, isosilybin ^60^, baicalein and scutellarein ^61^ has been described. Most of the reported studies were centered around simple flavanones or chalcones through enzymatic cascade reactions with the feeding of expensive phenyl precursors (phenylalanine, tyrosine, *p*-coumaric acid and caffeic acid etc), which limits our opportunity for scalable and low-cost production. Structural activity relationship (SAR) studies demonstrate the side chain modifications are highly correlated with flavonoid biological activities ^42, 62^. The hydroxylation of flavonoids improves their metabolic stability and membrane permeability, and enhances the solubility and antioxidant property ^63^. To date, there remains a significant challenge to produce highly hydroxylated flavonoids, due to our limited ability to functionally express plant P450 hydroxylases and the cytochrome P450 reductases ^64^. Oleaginous yeast is rich in membrane structure and subcellular compartments (i.e. lipid bodies, peroxisome, ER and oleosome), which provides the hydrophobic environment that is critical for regioselectivity and stereoselectivity in hydroxylation, glycosylation and prenylation of flavonoids ^65–68^. Developing a P450 expression platform in oleaginous yeast will enable us to access the vast majority of complex natural products and deliver robust microbial cell factories to meet the market demand.

To bridge this gap, we tested and assessed various plant-derived polyketide synthases, P450 monooxygenase/hydroxylases and cytochrome P450 reductases in *Y. lipolytica*, to diversify the structure of flavonoids. With naringenin, eriodyctiol and taxifolin as testing molecules, we characterized the catalytic efficiency of various plant enzymes, including tyrosine ammonia lyase (TAL), 4-coumaroyl-CoA ligase (4CL), chalcone synthase (CHS), chalcone isomerase (CHI), flavonone-3’-hydroxylase (F3’H), flavonol-3-hydroxyalse (F3H) and cytochrome P450 reductases. These plant-derived genes were coexpressed with the endogenous acetyl-CoA carboxylase (ACC1) and the pentafunctional AROM polypeptide (ARO1). Systematic pathway debottlenecking indicates that chalcone synthase, ACC1 and cytochrome P450 reductases are the rate-limiting steps for hydroxylated flavonoid production. Specifically, increasing PhCHS copy number and controlling culture pH elevated naringenin production up to 252.4 mg/L. Screening four cytochrome P450 reductases led us to identify that CrCPR derived from *Catharanthus roseus* is the most efficient electron shuttle to complete the hydroxylation reaction, despite that endogenous ylCPR1 (YALI0D04422g) displays similar function with relatively low efficiency. Further expression of the plant-derived P450 enzymes, including the flavanol-3’ hydroxylase (GhF3’H) from *Gerbera hybrid* led the engineered strain to produce about 110.5 mg/L of taxifolin and 134.2 mg/L of eriodictyol. This work set the foundation for us to engineer oleaginous yeast as chassis for cost-efficient production of flavonoids and hydroxylated flavonoids. The functional expression of plant-derived polyketide synthase, P450 monooxygenase and reductases will expand our capability to access nature’s biosynthetic potential for drug discovery and natural product manufacturing.

## Results and discussion

### Modular construction and characterization of flavonoid pathway in *Y. lipolytica*

The availability of intracellular malonyl-CoA was reported to be a rate-limiting step of flavonoid synthesis in many microorganisms ^69–71^. Considering the high acetyl-CoA and malonyl-CoA flux, we firstly reconstructed the synthetic pathway and validated the feasibility of using *Y. lipolytica* as the chassis to produce flavonoids. In addition, the cytochrome *c* P450 (CYP) flavonoid 3’-hydroxylase (F3’H) plays a critical role in oxidizing the phenyl ring and generating hydroxylated flavonoids ^72^. Based on the distribution of potential rate-limiting steps, we rationalized and partitioned the flavonoids pathway into two modules, the naringenin synthesis module (Module I) and the hydroxylation module (Module II) (Fig. 1). Module I contains essential precursor pathway to provide shikimic acid, malonyl-CoA and chalcone precursors; while Module II contains the cytochrome *c* P450 (CYP) flavonoid 3’-hydroxylase (F3’H) and cytochrome *c* P450 reductase (CPR). As a direct assessment of the module efficiency, we have established HPLC method to analyze naringenin, eriodyctiol and taxifolin (Fig. 2).

**Fig. 1.**
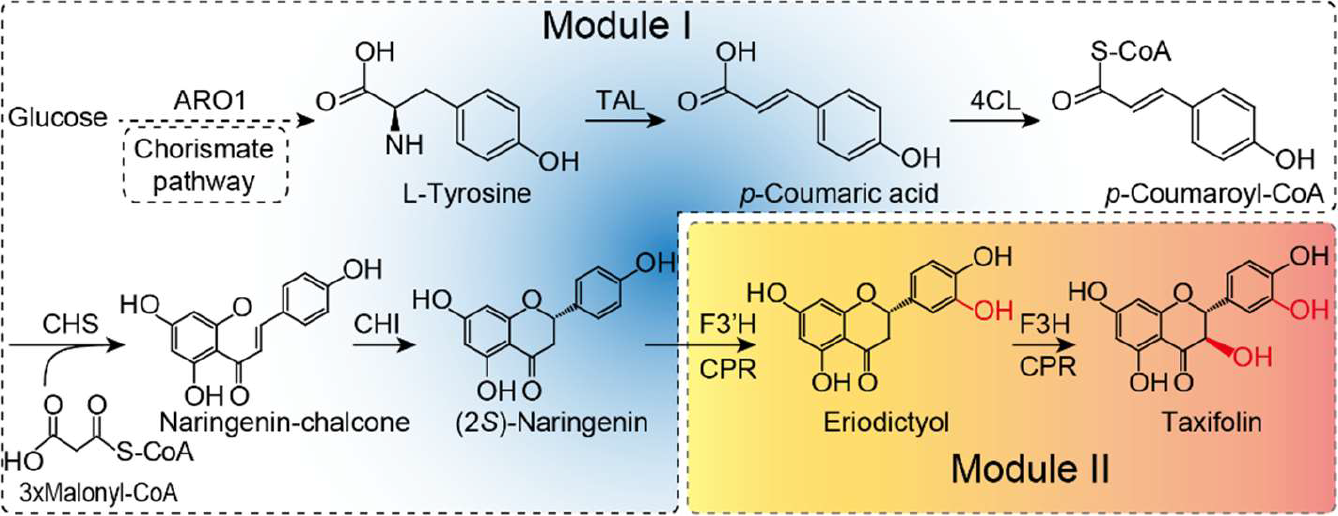
Modular strategy to optimize naringenin, eriodictyol and taxifolin pathways. Based on the reaction cascades, flavonoid pathway was partitioned into 2 modules, naringenin synthetic module (Module I) and hydroxylation module (Module II). Module I contains chorismate pathway and malonyl-CoA utilizing step, and Module II contains flavanone 3-hydroxyase and cytochrome c P450 reductases.

**Fig. 2.**
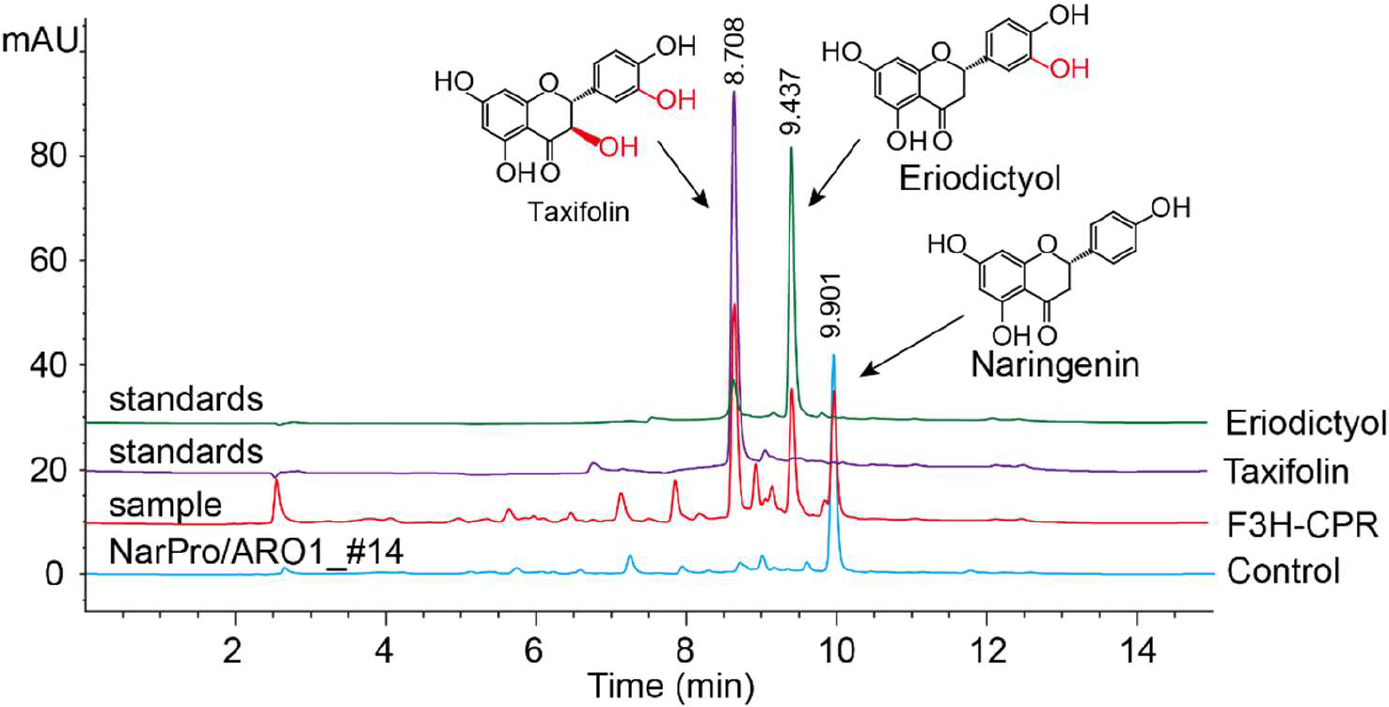
HPLC profile of naringenin, eriodictyol and taxifolin. Two hydroxylated flavonoid standards (taxifolin, purple and eriodictyol, green) were injected to HPLC. One naringenin-producing sample (blue) and one taxifolin-producing sample (red) are shown in the chromatogram.

Naringenin is the starting point for many flavonoid functionalization chemistry. We first constructed Module I in *Y. lipolytica* Po1f to synthesize chalcone and naringenin. Because genes from different plants have different specificity and activity, we selected two genes for each of the first three steps in Module I based on the sequence alignment of closely-related plant species. Pathways containing 4CL (*p*-coumaric acid-CoA ligase), CHS (chalcone synthase) and CHI (chalcone isomerase) were assembled in monocistronic forms by YaliBricks cloning platform ^34^. We observed that all eight constructs containing 4CL, CHS and CHI resulted in the synthesis of naringenin from *p*-coumaric acid, with production ranging from 10 mg/L to 21.5 mg/L (Fig. 3a). Interestingly, the three top producers (Fig. 3a) share the same source of chalcone synthase from *Petunia x hybrid*, indicating that chalcone synthase dictates the efficiency of Module I. To achieve *de novo* synthesis of naringenin, we further introduced tyrosine ammonia-lyase (RtTAL) from *Rhodotorula toruloides*, which has been reported to generate phenylpropanoid precursors from glucose ^47, 73^. With the overexpression of RtTAL, we detected *p*-coumaric acid as the direct de-amination product of tyrosine (Supplementary Fig. S1). By complementing the 4CL-CHS-CHI pathway, the resulted strain *Y. lipolytica* Po1f/T4SI produced 14.9 mg/L naringenin from glucose (Supplementary Fig. S1). These results validated the feasibility of using *Y. lipolytica* as chassis for *de novo* synthesis of naringenin.

**Fig. 3.**
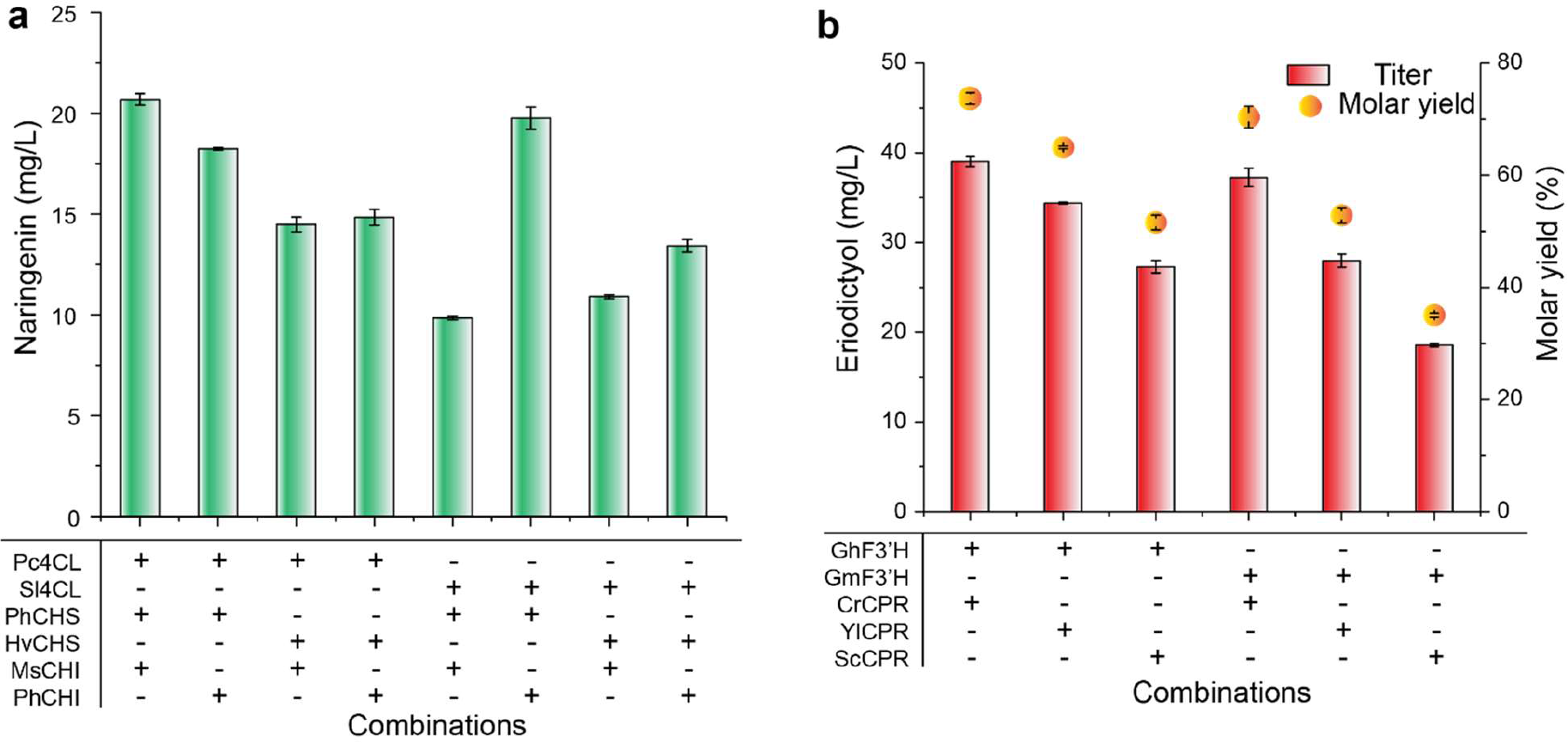
Screening of gene combinations for improving flavonoid production. (a) Screening of *4CH, CHS* and *CHI* genes from different pants for naringenin production. (b) Screening of *F3’H* and *CPR* genes from different organisms for eriodictyol production. Plant name and gene sources can be found in supplementary table S1.

There has been a number of reports that *Y. lipolytica* could selectively hydroxylate limonene to perillyl alcohol, perillaldehyde and perillic acids ^74, 75^, demonstrating the endogenous P450 monooxygenase and cytochrome P450 reductase is active enough to hydroxylate methyl group on monoterpenes. Our lab has demonstrated the functional expression of the P450 monooxygenase that selectively hydroxylates protodeoxyviolaceinic acid to protoviolaceinic acid, generating the greenish pigment in *Y. lipolytica* ^76^. On the basis of these results, we argue that *Y. lipolytic* could be an excellent platform for expression of plant P450 enzymes.

F3’H is the critical enzyme involved in the functional hydroxylation of flavonoids. Cytochrome *c* P450 reductase (CPR) is required for electron transfer from NADPH to CYP ^77^. We have chosen two plant-derived F3’Hs and three CPRs to evaluate which F3’H-CPR pairs could perform hydroxylation chemistry (Fig. 3b). All 6 combinations of F3’H-CPR pairs produced eriodictyol. We observed that strain Po1f/HR with overexpression of CrCPR (derived from *Catharanthus roseus*)^78^ coupled with GhF3’H (derived from *Gerbera hybrid*) or GmF3’H (derived from *Glycine max*), led to the highest eriodictyol production around 39 mg/L, with molar conversion yield up to 73.7% from naringenin (Fig. 3b). Interestingly, the two yeast-sourced CPRs, YlCPR from *Y. lipolytica* and ScCPR from *S. cerevisiae* S288c, also gave rise to eriodictyol, indicating the endogenous CPR is sufficient to shuttle electrons from NADPH to the active oxygen species, which is consistent with the findings reported by Leonard ^79^. We further constructed strain *Y. lipolytica* Po1f/HRH with the overexpression of flavanone 3-hydroxylase (SlF3H from *Solanum lycopersicum*) and detected about 26.0 mg/L taxifolin with a molar yield of 46.5% from naringenin (Fig. 2). In addition, *Y. lipolytica* endogenous CPR matched F3’H well, with comparable efficiency as CrCPR (Fig. 3b), which is consistent with previous report that *Y. lipolytica* was capable of performing P450-based biotransformation ^80, 81^. To achieve *de novo* synthesis of eriodictyol and taxifolin, we further complemented the RtTAL with GhF3’H and SlF3H, resulting in strains Po1f/T4SIHR and Po1f/T4SIHRH, respectively. When these strains were tested in shake flask cultures, we obtained 17.2 mg/L eriodictyol and 11.3 mg/L taxifolin from glucose, respectively. These results validated that *Y. lipolytica* will be an ideal chassis to functionally express plant P450 enzymes and produce hydroxylated flavonoids.

### Tuning gene-copy number to remove pathway bottlenecks

The balance of metabolic flux and mitigation of metabolic burden is a vital factor for optimizing metabolite production in microorganisms ^9, 53, 82^. Introduction of large gene cluster may result in the host strain losing cellular fitness when the expression of heterologous proteins exceeds the carrying capacity of the system. For example, metabolic flux improvement by overexpression of upstream pathways may not be accommodated by downstream pathways ^82^; intermediate accumulation or depletion may reduce cell viability ^83^; and overexpressed gene clusters may overload the cell and elicit cellular stress response ^84, 85^. We next attempted to probe the rate-limiting steps in Module I and Module II by gradually increasing gene copy number of the genes involved. Gene copy number of each enzymatic step was individually tuned by using YaliBrick assembly platform ^34^. Naringenin production increased by 2.64-fold when the gene copy number for chalcone synthase (PhCHS) increased from one to five (Fig. 4a), indicating that CHS is the rate-limiting step in Module I. Increasing the gene copy number of other metabolic genes (RtTAL and Pc4CL) did not have obvious effect on naringenin titer, while increasing the gene copy number of MsCHI decreased naringenin titer by 34.4% (Fig. 4a). We determined that the optimal gene copy number for *PhCHS* is 5, as naringenin production was only marginally increased when the gene copy number was changed from 4 to 5. As larger plasmid may cause genetic instability, we did not further increase the copy number of *PhCHS*.

**Fig. 4.**
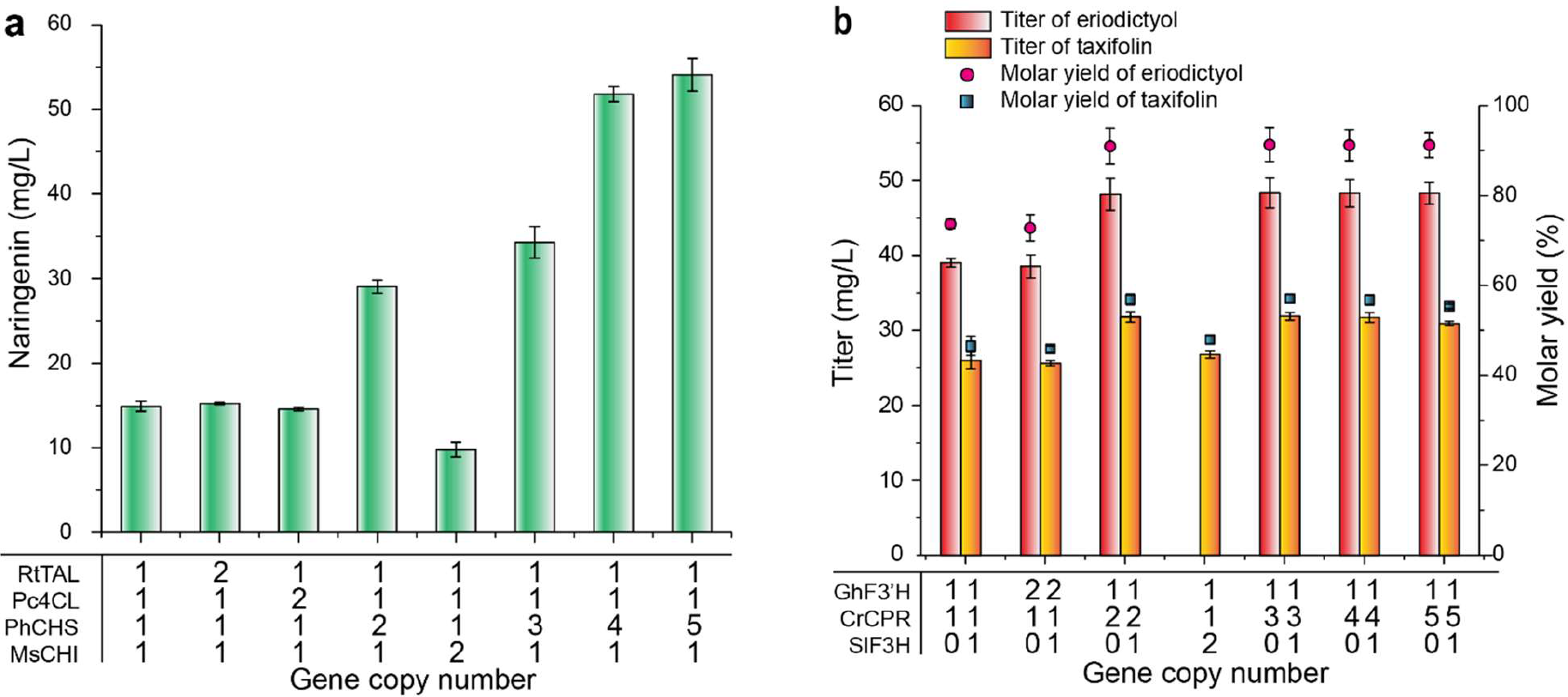
Overcoming rate-limiting steps by tuning gene copy numbers. Rate-limiting steps were determined by gradually increasing gene copy number of each step. Numbers refer to gene copy numbers. (a) Rate-limiting step analysis and optimization of module I to improve naringenin production. (b) Rate-limiting step analysis and optimization of module II to improve eriodictyol and taxifolin production. The molar yield was calculated using 50 mg/L naringenin as feeding substrate. The number 0 refers to the module that does not contain the respective gene.

In Module II, increasing one copy number of *CrCPR* resulted in eriodictyol and taxifolin titers increasing by 26.8% and 22.3%, reaching 48.1 mg/L and 31.8 mg/L (Fig. 4b), respectively. Increasing the copy number of *SlF3H* and *GhF3’H* did not have obvious effect on eriodictyol or taxifolin production (Fig. 4b), indicating that CPR is the rate-limiting step in Module II. Eriodictyol and taxifolin titers remained stable, when the gene copy number for *CrCPR* was increased from 2 to 5, suggesting that the optimal ratio of F3’H to its reductase *CrCPR* is 1:2, which is consistent with previous report ^86^. The naringenin-to-eriodictyol and taxifolin conversion ratio reached 90.5% and 56.8%, respectively (Fig. 4b), under the optimal F3’H-CPR ratios. To achieve *de novo* synthesis of eriodictyol and taxifolin, we complemented the eriodictyol and taxifolin pathways with the RtTAL pathway. The resulting strains Po1f/T4S_x5_IHR_x2_ and Po1f/T4S_x5_IHR_x2_H produced 28.9 mg/L eriodictyol and 25.2 mg/L taxifolin from glucose, respectively. These titers are 68.0% and 123.0% higher than the control strains Po1f/T4SIHR and Po1f/T4SIHRH. These results confirmed that tuning gene copy numbers will be a critical step to remove pathway bottlenecks and achieve metabolic balance in genetically modified cell factories, in particularly, oleaginous yeast for flavonoids production.

### Improving flavonoid production by enhancing precursor synthesis

We next sought to investigate the upstream shikimic acid and malonyl-CoA pathways to further improve flavonoids production. By supplementing 100 mg/L L-tyrosine with the strain Po1f/T4SI, we observed that naringenin production was increased by 33.6% with glucose as sole carbon source, indicating that upstream shikimic acid pathway is a bottleneck for naringenin synthesis in *Y. lipolytica*. We then overexpressed the pentafunctional polypeptides arom protein ARO1, which catalyzes steps 2 through 6 in the biosynthesis of chorismate, to boost the precursor for L-tyrosine synthesis ^87^. YALI0F12639g (YlARO1) is a *Y. lipolytica* homologue of S. cerevisiae ARO1 ^88^, and the DNA sequence for YlARO1 is composed of 1 intron and 2 exons, encoding a 1556-aa protein. To mitigate unintended mRNA splicing and transcriptional regulation, we removed the internal intron for YlARO1. When this *YlARO1* gene was overexpressed in strain Po1f/T4S_x5_I with optimal Module I settings, naringenin production was increased to 81.6 mg/L, a 50.9% increased compared to the parental strain (Fig. 5a). When we combined Module I with Module II, the resulting strains Po1f/AT4S_x5_IHR_x2_ and Po1f/AT4S_x5_IHR_x2_H produced 40.1 mg/L eriodictyol and 33.4 mg/L taxifolin, which is 38.8% and 32.5% higher than that of the control strains Po1f/T4S_x5_IHR_x2_ and Po1f/T4S_x5_IHR_x2_H, respectively (Fig. 5b).

**Fig. 5.**
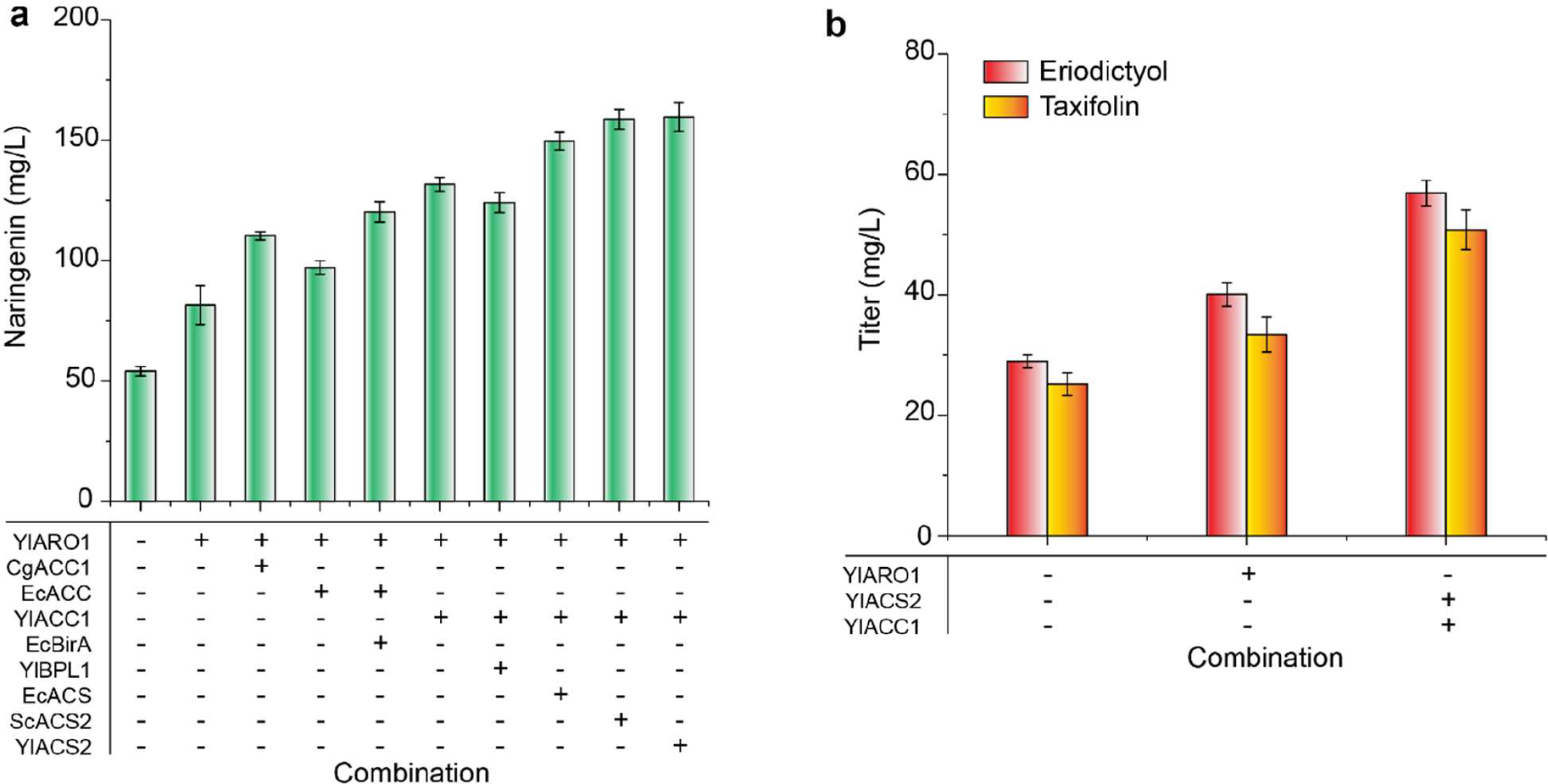
Improving naringenin, eriodictyol, and taxifolin production by enhancing precursor synthesis. (a) Identification of possible rate-limiting steps by overexpression of chorismate pathway (ARO1), malonyl-CoA pathway (ACC) and acetyl-CoA pathway (ACS). The related genes were overexpressed in strains Po1f/T4S_x5_I. (b) Effects of improving malonyl-CoA and chorismate synthesis on eriodictyol and taxifolin production. For eriodictyol production, the related genes were overexpressed in Po1f/T4S_x5_IHR_x2_. For taxifolin production, the related genes were overexpressed in Po1f/T4S_x5_IHR_x2_H. + referred to the presence of gene overexpression. – referred to the absence of gene overexpression.

Acetyl-CoA and malonyl-CoA are shared precursors for both lipids and flavonoid pathway ^89, 90^. However, malonyl-CoA is primarily used to synthesize lipids cell membrane and support cell growth, leaving only a small amount of acetyl-CoA and malonyl-CoA for heterologous production. To mitigate this competition, it is desirable to redirect the acetyl/malonyl-CoA flux from lipid pathway to flavonoid pathway. Acetyl-CoA carboxylase (ACC) converts acetyl-CoA to malonyl-CoA, which is the first committed step in both lipid and flavonoids biosynthesis ^69^. In order to enhance intracellular malonyl-CoA synthesis, we screened and tested different ACCs from three sources, including gram-positive bacteria *Corynebacterium glutamicum* ATCC 13032 (*CgACC*), gram-negative bacteria *Escherichia coli* MG1655 (Ec_*accABCD*), and *Y. lipolytica* (*YlACC1*, GRYC ID: YALI0C11407g) ^91, 92^. Biotin-apoprotein ligase modifies ACC by covalently attaching biotin, which is essential for ACC activity ^93^. *EcBirA* and YlBPL1 (YALI0E30591g) are *E. coli* and *Y. lipolytica* homologues of biotin-apoprotein ligase, respectively ^94, 95^. Genes encoding CgACC, Ec_accABCD, and YlACC1, together with their biotin-apoprotein ligases were introduced to the naringenin-producing strain. All three ACCs could lead to substantial improvement in naringenin production (Fig. 5a), with YlACC1 demonstrating most obvious effect. For example, overexpression of *YlACC1* in Po1f/AT4S_x5_I improved naringenin titer by 61.4%, reaching 131.7 mg/L (Fig. 5a). The coupling of *Ec_accABCD* with *EcBirA* also resulted in naringenin production increasing by 22% compared with the strain without *EcBirA* overexpression, indicating the essential role of biotinylation in bacterial ACC activity. This is the first report that *Ec*ACC could be functionally expressed in oleaginous species. Unlike the bacterial ACC, co-expression of *YlACC1* and *YlBPL1* resulted in decreased naringenin production (Fig. 5a). This might indicate the endogenous biotin-apoprotein ligase (YlBPL1) is sufficient to biotinylate ylACC1 in *Y. lipolytica*.

We observed that pH value dropped dramatically during the fermentation process (i.e. pH below 3.5 at the end of flask cultivation), and this could be largely ascribed to the overflown metabolism of TCA cycle and respiration ^20^. It was recently discovered that acetate secretion was associated with the CoA-transfer reaction between acetyl-CoA and succinate in *Y. lipolytica*, encoded by a mitochondrial enzyme ylACH1 (YALI0E30965) ^23^. To recycle acetate, we next sought to overexpress acetyl-CoA synthetases and convert acetate to acetyl-CoA. We tested three acetyl-CoA synthetases from *E. coli*, *S. cerevisiae* and *Y. lipolytica* (Fig. 5a). The native version *YlACS2* demonstrate better effect to recycle acetate. To test the combinatory effects of enhancing chorismate and acetyl/malonyl-CoA precursors, we overexpressed YlARO1 along with YlACS2-YlACC1 in strain Po1f/T4S_x5_I. The resulting strain produced 149.5 mg/L naringenin, which was 176.3% higher than the titer of the parental strain (Po1f/T4S_x5_I). We further applied the same strategy to Module II and tested whether overexpression of ARO1, ACC1 and ACS would benefit the accumulation of hydroxylated flavonoids. Overexpression of *YlARO1* increased eriodictyol and taxifolin production by 38.8% and 32.5%, yielding 40.1 mg/L and 33.4 mg/L (Fig. 5b), respectively. Overexpressing *YlACS2* and *YlACC1* further increased eriodictyol and taxifolin titers by 41.9% and 52.1%, reaching 56.9 mg/L and 50.8 mg/L, respectively (Fig. 5b). Due to the large size of the plasmid construct (more than 40 kb), we did not further pursue the synergistic effect of ARO1, ACC1 and ACS2 in the current work. These results indicated that manipulation of acetyl-CoA, malonyl-CoA and chorismate pathway was critical to improve flavonoid production in *Y. lipolytica*.

### Boosting flavonoid production by bioprocess optimization

The C/N ratio is an important factor for regulating the acetyl-CoA and NADPH fluxes in *Yarrowia lipolytica* ^33^. It has been reported that nitrogen starvation triggers the repression of TCA cycle and induces lipogenesis in oleaginous species ^96, 97^. It was recently discovered that C/N ratio dynamically regulates lipogenic promoter activity in *Y. lipolytica* ^33^. In this study, C/N ratio was optimized in two patterns to improve flavonoid synthesis, by either adjusting the amount of nitrogen source (ammonia sulfate) or carbon source (glucose). The results showed that altering (NH_4_)_2_SO_4_ content did not have obvious effect on naringenin titer. Slightly higher naringenin titer was achieved at higher C/N ratio (C/N=120) (Supplementary Fig. S2a). On the contrary, it was clearly shown that higher C/N ratio was advantageous to improving naringenin titer by increasing the level of glucose. Specifically, naringenin titer was increased about 56% when the C/N was altered from 40 to 160 (Supplementary Fig. S2b). Since glucose is the direct precursor for chorismate and malonyl-CoA, it indicates that there is still much space to further improve the precursor flux in *Y. lipolytica*.

In order to produce flavonoids with inexpensive YPD (yeast extract, peptone and dextrose) medium, we integrated the optimized pathways into *Y. lipolytica* Po1f genome with our recently developed integration methods ^98^. The best-performing strains NarPro/ASC, ErioPro, and TaxiPro produced 71.2 mg/L naringenin, 54.2 mg/L eriodictyol, and 48.1 mg/L taxifolin in YPD medium, respectively ^40^. We observed that the pH dropped to 3.2 at the end of the fermentation in YPD, possibly due to the accumulation of various organic acids ^20^. We next sought to buffer the media pH by using either phosphate buffer saline (PBS) or calcium carbonate (CaCO_3_). Supplementation of 4% CaCO_3_ maintained stable pH and improved naringenin titer by 31.2%, reaching 138.1 mg/L at 144 h, while PBS buffer did not have obvious effect compared with the control (Supplementary Fig. S4). We also analyzed the combinatory effects of inhibiting fatty acid synthesis by adding cerulenin ^99^, maintaining stable pH, and supplying sodium acetate. The fermentation time course showed that we could achieve steady improvement in naringenin production (Supplementary Fig. S5). For example, supplementation of 1 mg/L cerulenin with 40 g/L CaCO_3_ further improved naringenin titer by 31.2%, (Supplementary Fig. S5c). However, supplying 5 mM NaAc did not result in further increase in naringenin production (Supplementary Fig. S5d). By intermittently feeding glucose after 48 hours, the chromosomally-integrated strain produced 252.4 mg/L naringenin under optimal conditions (Fig. 6a). Likewise, we tested the eriodictyol and taxifolin production in YPD medium by buffering media pH with 40 g/L CaCO_3_ and inhibiting fatty acid synthesis with 1 mg/L cerulenin. We observed that strains ErioPro and TaxiPro produced 95.5 mg/L eriodictyol and 79.1 mg/L taxifolin in 144 h, which were 76.2% and 64.4% higher than that without CaCO_3_ and cerulenin. ErioPro and TaxiPro produced 134.2 mg/L eriodictyol and 110.5 mg/L taxifolin at the end of the fermentation process (Fig. 6b). These results indicate that *Y. lipolytica* is an ideal platform to functionally express plant-derived P450 enzymes. By optimizing the bioprocess, we could substantially improve the titer of naringenin, eriodictyol and taxifolin in metabolically-engineered oleaginous yeast species.

**Fig. 6.**
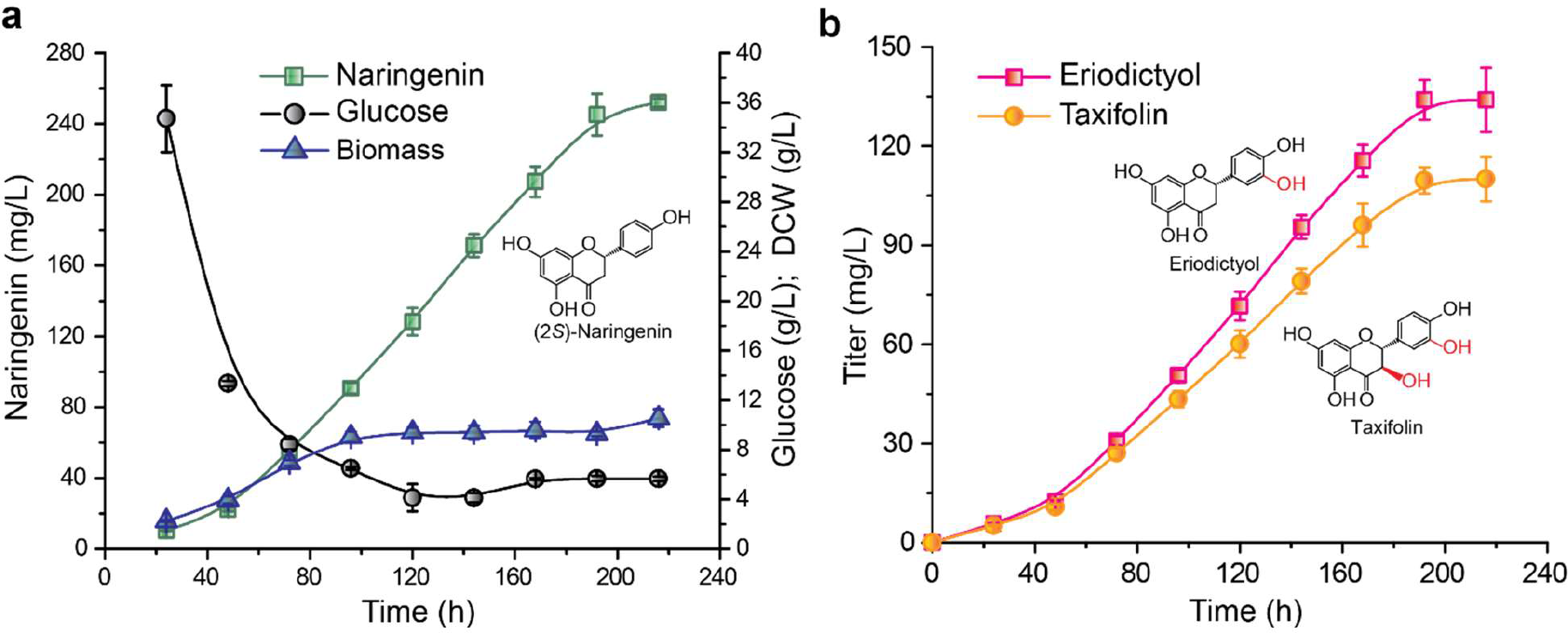
Naringenin, eriodictyol and taxifolin production under the optimal conditions. (a). Naringenin production; (b) Eriodictyol and taxifolin production. The engineered strains were cultivated in fed-batch fermentation and buffered with 40 g/L CaCO_3_. A final concentration of 1 mg/L cerulenin was supplemented at 48 h to inhibit fatty acid synthesis.

## Conclusions

The heterologous production of hydroxylated flavonoids remains a challenging task; and only limited successful pathway engineering endeavors have been reported to date. Oleaginous yeast is rich in lipid and internal membrane structures, which provides the hydrophobic lipid environment and spatially-organized organelles that are critical for plant P450 enzyme functionality. In this report, we validated that *Y. lipolytica* is a superior platform for heterologous production of high value flavonoids and hydroxylated flavonoids. By modular construction and characterization of various genes involved in plant flavonoid biosynthesis, we determined that chalcone synthase (CHS), flavanone 3-hydroxylase (F3H) and cytochrome c P450 reductase (CrCPR) were the critical steps to engineer flavonoid production in *Y. lipolytica*. Coupling with the upstream amino acid degradation pathway (tyrosine ammonia lyase from *Rhodotorula glutinis*), for the first time, we achieved *de novo* production of naringenin, eriodictyol and taxifolin from glucose in *Y. lipolytica*. By using a modular cloning platform to assemble multiple genetic constructs, we further determined the optimal gene copy ratio for CHS, F3H and CrCPR to cooperatively improve flavonoids and hydroxylated flavonoids production. We then unleashed the metabolic potential of the yeast host by screening and testing a number of precursor pathways, including the acetyl-CoA synthetase, acetyl-CoA carboxylase and chorismate pathway (the pentafunctional AROM polypeptide ARO1). Coupled with the optimized chalcone synthase module and the hydroxylation module, our engineering strategies synergistically removed pathway bottlenecks and led to a 15.8-fold, 6.9-fold and 8.8-fold improvement in naringenin, eriodictyol and taxifolin production, respectively. Collectively, these findings demonstrate our abilities to harness oleaginous yeast as microbial workhorse to expand nature’s biosynthetic potential, which allows us to produce complex natural products from cheap feedstocks.

## Materials and methods

### Genes, plasmids, and strains

Genes encoding *Rhodotorula R. toruloides* tyrosine ammonia lyase (*RtTAL*), *Petroselinum crispum* (parsley) 4-coumarate-CoA ligase (*Pc4CL*), *Petunia x hybrid* chalcone synthase (*PhCHS*), *Medicago sativa* chalcone isomerase (*MsCHI*), *Escherichia coli* acetyl-CoA synthetase (*EcACS*), and *Corynebacterium glutamicum* ATCC 13032 acetyl-CoA carboxylase (*CgACC*) were frozen stocks of our laboratory. Genes encoding *Solanum lycopersicum* 4-coumarate-CoA ligase (*Sl4CL*), *Hordeum vulgare* chalcone synthase (*HvCHS2*), *Petunia x hybrid* chalcone isomerase (*PhCHI*), *Gerbera hybrid* flavonoid 3’-hydroxylase (*GhF3’H*), *Glycine max* flavonoid 3’-hydroxylase (*GmF3’H*), *Catharanthus roseus* cytochrome P450 reductase (*CrCPR*), and *Solanum lycopersicum* flavanone 3-hydroxylase (*SlF3H*) were optimized and synthesized by GenScript (Nanjing, China). Genes encoding *Yarrowia lipolytica* pentafunctional arom protein (*YlARO1*), *Yarrowia lipolytica* cytochrome P450 reductase (*YlCPR*), *Yarrowia lipolytica* acetyl-CoA synthetase (*YlACS2*) were amplified from *Yarrowia lipolytica* Po1f genomic DNA by PCR. *Saccharomyces cerevisiae* cytochrome P450 reductase (*ScCPR1*) and *Saccharomyces cerevisiae* acetyl-CoA synthetase (*ScACS2*) were amplified from *Saccharomyces cerevisiae* genomic DNA by PCR. Genes used in this project were listed in Supplementary Table S1.

Plasmid pYLXP’ was a stock of our laboratory ^89^. Plasmid pYLXP’2 was constructed by replacing *LEU2* marker with *URA3* marker. Both pYLXP’ and pYLXP’2 were YaliBrick plasmids and used for flavonoid pathway construction ^34^. *Escherichia coli* (*E. coli*) NEB 5α was used for plasmid construction, propagation, and maintenance. *Yarrowia lipolytica* (*Y. lipolytica*) Po1f (ATCC MYA-2613, MATA ura3-302 leu2-270 xpr2-322 axp2-deltaNU49 XPR2::SUC2) was used as the chassis to construct flavonoid pathways.

To achieve *de novo* synthesis of eriodictyol and taxifolin, we transformed pYLXP’2-GhF3’H-CrCPR and pYLXP’2-GhF3’H-CrCPR-SlF3H into Po1f/T4SI, resulting in Po1f/T4SIHR and Po1f/T4SIHRH, respectively. The strain containing 5 copies of *PhCHS* was named as Po1f/T4S_x5_I. We chose to use the plasmids pYLXP’2-HR_x2_ and pYLXP’2-HR_x2_H, which contain 2 copies of *CrCPR*, to construct eriodictyol and taxifolin pathways. To achieve *de novo* synthesis of eriodictyol and taxifolin, strains Po1f/T4S_x5_IHR_x2_ and Po1f/T4S_x5_IHR_x2_H were constructed by transforming plasmids pYLXP’2-HR_x2_ and pYLXP’2-HR_x2_H into strain Po1f/T4S_x5_I, respectively. We over-expressed YlARO1 along with YlACS2-YlACC1 in strain Po1f/T4S_x5_I, and name the new strain as Po1f/AT4S_x5_I-YlACS2-YlACC1. By introducing Module II into strains Po1f/AT4S_x5_I and Po1f/AT4S_x5_I-YlACS2-YlACC1, we obtained eriodictyol producing strains Po1f/AT4S_x5_IHR_x2_ and Po1f/AT4S_x5_IHR_x2_-YlACS2-YlACC1 and taxifolin producing strains Po1f/AT4S_x5_IHR_x2_H and Po1f/AT4S_x5_IHR_x2_H-YlACS2-YlACC1. Strains constructed in this project were listed in Supplementary Table S2.

### Pathway construction

Genes *RtTAL, Pc4CL, PhCHS, MsCHI, EcACS, CgACC, YlCPR, YlACS2, ScCPR1*, and *ScACS2* were amplified using respective primers listed in Supplementary Table S3. The PCR product was assembled with *SnaBI* digested pYLXP’ or pYLXP’2 using Gibson Assembly method. *YlARO1* is composed of 2 extrons and 1 intron. The extrons were amplified by using primer pairs ARO1_up F/ARO1_up R and ARO1_down F/ARO1_down R respectively. The resulting PCR products were assembled with *SnaBI* digested pYLXP’ to yield pYLXP’-ARO1, removing the intron sequence. For gene expression, the start codon was removed and a nucleic acid sequence “TAACCGCAG” was added at the upstream of coding gene to complete the intron ^34^.

The YaliBrick method was used to assemble the synthetic pathways ^34^. pYLXP’ derived plasmids were used to assemble the pathways of Module I, while pYLXP’2 derived plasmids were used to assemble the pathways of Module II and ACS and ACC. Generally, the donor plasmids were digested with *Avr*II/*Sal*I, and the destination plasmids were digested with *Nhe*I/*Sal*I. The resulting plasmids containing monocistronic configurations were obtained by T4 ligation. For the assemble of genes containing any of these isocaudomers, other isocaudomers were used. Specifically, the donor plasmid pYLXP’-YlARO1 was digested with HpaI/NheI, and the destination plasmid pYLXP’-T4S_x5_I was digested with *Hpa*I/*Avr*II. The resulting plasmid pYLXP’-AT4S_x5_I was obtained by inserting *YlARO1* into pYLXP’-T4S_x5_I using T4 ligation. The donor plasmid pYLXP’2-HR_x2_H was digested with *Cla*I/*Nhe*I, and the destination plasmid pYLXP’2-ScACS2-YlACC1 was digested with *Cla*I/*Avr*II. The resulting plasmid pYLXP’2-HR_x2_H-ScACS2-YlACC1 was obtained by inserting genes GhF3’H-CrCPR_x2_-SlF3H into pYLXP’2-ScACS2-YlACC1 using T4 ligation. Plasmids pYLXP’2-YlACC1, pYLXP’2-EcACCABCD-EcBirA, and pYLXP’2-YlBPL1 were frozen stocks of our laboratory^34^. Plasmids used in this paper were listed in Supplementary Table S4.

### Yeast transformation and screening

The lithium acetate (LiAc) method was used for the transformation. *Y. lipolytica* was cultured on YPD plate at 30°C for 16-22 h. The transformation solution was prepared as follows: 90 μL 50% PEG4000, 5 μL 2 M LiAc, 5 μL boiled single strand DNA (salmon sperm, denatured), and 200-500 ng plasmid DNA. The transformation solution was mixed well by vortexing before use. Next, the yeast was transferred to the transformation solution, and mixed well by vortexing for at least 10 seconds. The transformation mixtures were then incubated at 30°C for 30-45 min. The transformation mixture was then vortexed for 15 seconds every 10 minutes, followed by an additional 10 min heat shock at 39°C to increase transformation efficiency. For the transformation of pYLXP’ and derivative plasmids, the mixture was plated on leucine drop-out complete synthetic media (CSM-Leu). For the transformation of pYLXP’2 and derivative plasmids, the mixture was plated on uracil drop-out complete synthetic media (CSM-Ura). For the transformation of both plasmids, the mixture was plated on leucine and uracil drop-out complete synthetic media (CSM-Leu-Ura). Strains NarPro/ASC, ErioPro, and TaxiPro were constructed in previous work ^40^. Strains used in this paper were listed in Strains constructed in this project were listed in Supplementary Table S2.

### Cultivation and pH control

The seed was cultured in regular leucine, or uracil, or leucine and uracil drop-out complete synthetic media (CSM-Leu, or CSM-Ura, or CSM-Leu-Ura) at 30°C for 2 days. The seed culture was inoculated to 25 mL nitrogen-limited media (C/N = 80) to a final concentration of 2% (v/v). The fermentation was carried out in 250 mL shak flask at 30°C 220 rpm. C/N ratio was optimized by two patterns: i) fixing glucose content (40 g/L) and altering (NH_4_)_2_SO_4_ content; ii) fixing (NH_4_)_2_SO_4_ content (0.73348 g/L) and altering glucose content. To analyze the effect of cerulenin, oleic acid, and sodium acetate (NaAc) on flavonoid synthesis, a final concentration of 5 g/L oleic acid or 1 mM NaAc was added at the starting point, while a final concentration of 1 mg/L cerulenin was added at 48 h. To buffer the acidity, 20 mM phosphate buffer saline (PBS, Na_2_HPO_4_-NaH_2_PO_4_) or 40 g/L CaCO_3_ was used respectively. In the fed-batch fermentation, the starting glucose concentration was 40 g/L, and a final concentration of 10 g/L glucose was added every 24 h from 48 h.

### Analytical methods

Samples were taken at 144 h. In the fed-batch fermentation, samples were taken every 24 h. For naringenin, eriodictyol, and taxifolin analysis, samples were diluted in methanol; whole for glucose analysis, samples were diluted in H_2_O. Samples were shaken with glass beads to release the metabolites for analysis. Naringenin, eriodictyol, taxifolin, and glucose were analyzed using Agilent HPLC 1220 as previously described 40.

## Acknowledgements

This work was supported by the Cellular & Biochem Engineering Program of the National Science Foundation under grant no.1805139, the National Natural Science Foundation of China (31670095, 31770097), the Fundamental Research Funds for the Central Universities (JUSRP51701A), the Distinguished Professor Project of Jiangsu Province. The authors would also like to acknowledge the Department of Chemical, Biochemical and Environmental Engineering at University of Maryland Baltimore County for funding support. YL would like to thank the China Scholarship Council for funding support.

## Author contributions

PX and JZ conceived the topic. YL performed genetic engineering and fermentation experiments. YL and PX wrote the manuscript. JZ and MK revised the manuscript.

