## Supplemental Tables and figures for "Optimizing oleaginous yeast cell factories for flavonoids and hydroxylated flavonoids biosynthesis"

**Table S1. Genes used in this paper.**

| Gene | Origin | Database and accession number | Optimization | References |
|---|---|---|---|---|
| *RtTAL* | *Rhodotorula toruloides* | NCBI: AAA33883 | Yes | (*1*) |
| *Pc4CL* | *Petroselinum crispum* (parsley) | UniProtKB: P14912 | Yes | (*2*) |
| *Sl4CL* | *Solanum lycopersicum* | NCBI: NP_001333770 | Yes | (*3*) |
| *PhCHS* | *Petunia x hybrid* | NCBI: AAF60297 | Yes | (*2*) |
| *HvCHS2* | *Hordeum vulgare* | NCBI: Q96562 | Yes | (*4*) |
| *MsCHI* | *Medicago sativa* | NCBI: P28012 | Yes | (*2*) |
| *PhCHI* | *Petunia x hybrid* | NCBI: P11650 | Yes | (*5*) |
| *GhF3'H* | *Gerbera hybrid* | UniProtKB: Q38L00 | Yes | (*6*) |
| *GmF3'H* | *Glycine max* | NCBI: NP_001237015 | Yes | (*7*) |
| *CrCPR* | *Catharanthus roseus* | UniProtKB: Q05001 | Yes | (*8*) |
| *SlF3H* | *Solanum lycopersicum* | NCBI: NP_001316412 | Yes | (*9*) |
| *YlCPR1* | *Yarrowia lipolytica* Po1f | GRYC[1]: YALI0D04422g1_1 | No | |
| *ScCPR1* | *Saccharomyces cerevisiae* S288c | GRYC: YHR042W1_1 | No | |
| *YlACS2* | *Yarrowia lipolytica* Po1f | GRYC: YALI0F05962g1_1 | No | |
| *ScACS2* | *Saccharomyces cerevisiae* S288c | GRYC: YLR153C1_1 | No | |
| *EcACS* | *Escherichia coli* str. K12 substr. MG1655 | NCBI: AVI54216 | No | (*6*) |
| *YlACC1* | *Yarrowia lipolytica* Po1f | GRYC: YALI0C11407g1_1 | No | |
| *CgACC*[2] | *Corynebacterium glutamicum* ATCC 13032 | NCBI: NP_599932 and AUI00288 | No | (*6*) |

| | | | | |
|---|---|---|---|---|
| *YlARO1* | *Yarrowia lipolytica* Po1f | GRYC: YALI0F12639g1_1 | No | (*10*) |
| *EcACCABCD* | *Escherichia coli* DH 5α | | No | (*11*) |
| *EcBirA* | *Escherichia coli* DH 5α | | No | (*11*) |
| *YlBPL1* | *Yarrowia lipolytica* Po1f | GRYC: YALI0E30591g | No | |

(1) GRYC refers to the genome resources for yeast chromosomes (http://gryc.inra.fr/index.php?page=home).

(2) *CgACC* is a chimeric gene, composed of *accBC* and *dtsR1*.

**Table S2. Strains constructed in this paper.**

| Strain | Annotation | References |
|---|---|---|
| Po1f/Pc4CL-PhCHS-MsCHI | Po1f containing pYLXP'-Pc4CL-PhCHS-MsCHI, used for the optimization of Module I | This work |
| Po1f/Pc4CL-PhCHS-PhCHI | Po1f containing pYLXP'-Pc4CL-PhCHS-PhCHI, used for the optimization of Module I | This work |
| Po1f/Pc4CL-HvCHS2-MsCHI | Po1f containing pYLXP'-Pc4CL-HvCHS2-MsCHI, used for the optimization of Module I | This work |
| Po1f/Pc4CL-HvCHS2-PhCHI | Po1f containing pYLXP'-Pc4CL-HvCHS2-PhCHI, used for the optimization of Module I | This work |
| Po1f/Sl4CL-PhCHS-MsCHI | Po1f containing pYLXP'-Sl4CL-PhCHS-MsCHI, used for the optimization of Module I | This work |
| Po1f/Sl4CL-PhCHS-PhCHI | Po1f containing pYLXP'-Sl4CL-PhCHS-PhCHI, used for the optimization of Module I | This work |
| Po1f/Sl4CL-HvCHS2-MsCHI | Po1f containing pYLXP'-Sl4CL-HvCHS2-MsCHI, used for the optimization of Module I | This work |
| Po1f/Sl4CL-HvCHS2-PhCHI | Po1f containing pYLXP'-Sl4CL-HvCHS2-PhCHI, used for the optimization of Module I | This work |
| Po1f/GhF3'H-CrCPR | Po1f containing pYLXP'2-GhF3'H-CrCPR, used for the optimization of Module II | This work |
| Po1f/GhF3'H-YlCPR | Po1f containing pYLXP'2-GhF3'H-YlCPR, used for the optimization of Module II | This work |
| Po1f/GhF3'H-ScCPR | Po1f containing pYLXP'2-GhF3'H-ScCPR, used for the optimization of Module II | This work |
| Po1f/GmF3'H-CrCPR | Po1f containing pYLXP'2-GmF3'H-CrCPR, used for the optimization of Module II | This work |
| Po1f/GmF3'H-YlCPR | Po1f containing pYLXP'2-GmF3'H-YlCPR, used for the optimization of Module II | This work |
| Po1f/GmF3'H-ScCPR | Po1f containing pYLXP'2-GmF3'H-ScCPR, used for the optimization of Module II | This work |
| Po1f/T4SI | Po1f containing pYLXP'-T4SI, used for rate-limiting step analysis and optimization of Module I | This work |
| Po1f/$T_{x2}$4SI | Po1f containing pYLXP'-$T_{x2}$4SI, used for rate-limiting step analysis and optimization of Module I | This work |
| Po1f/$T4_{x2}$SI | Po1f containing pYLXP'-$T4_{x2}$SI, used for rate-limiting | This work |

| | | |
|---|---|---|
| | step analysis and optimization of Module I | |
| Po1f/T4S$_{x2}$I | Po1f containing pYLXP'-T4S$_{x2}$I, used for rate-limiting step analysis and optimization of Module I | This work |
| Po1f/T4SI$_{x2}$ | Po1f containing pYLXP'-T4SI$_{x2}$, used for rate-limiting step analysis and optimization of Module I | This work |
| Po1f/T4S$_{x3}$I | Po1f containing pYLXP'-T4S$_{x3}$I, used for the optimization of Module I | This work |
| Po1f/T4S$_{x4}$I | Po1f containing pYLXP'-T4S$_{x4}$I, used for the optimization of Module I | This work |
| Po1f/T4S$_{x5}$I | Po1f containing pYLXP'-T4S$_{x5}$I, used for the optimization of Module I | This work |
| Po1f/HRH | Po1f containing pYLXP'2-HRH, used for the rate-limiting step analysis and optimization of Module II | This work |
| Po1f/H$_{x2}$RH | Po1f containing pYLXP'2-H$_{x2}$RH, used for the rate-limiting step analysis and optimization of Module II | This work |
| Po1f/HR$_{x2}$H | Po1f containing pYLXP'2-HR$_{x2}$H, used for the rate-limiting step analysis and optimization of Module II | This work |
| Po1f/HRH$_{x2}$ | Po1f containing pYLXP'2-HRH$_{x2}$, used for the rate-limiting step analysis and optimization of Module II | This work |
| Po1f/HR$_{x3}$H | Po1f containing pYLXP'2-HR$_{x3}$H, used for the optimization of Module II | This work |
| Po1f/HR$_{x4}$H | Po1f containing pYLXP'2-HR$_{x4}$H, used for the optimization of Module II | This work |
| Po1f/HR$_{x5}$H | Po1f containing pYLXP'2-HR$_{x5}$H, used for the optimization of Module II | This work |
| Po1f/AT4S$_{x5}$I | Po1f containing pYLXP'-AT4S$_{x5}$I, used for enhancing the synthesis of chorismate (L-tyrosine) | This work |
| Po1f/AT4S$_{x5}$I-CgACC1 | Po1f/AT4S$_{x5}$I containing pYLXP'2-CgACC1, used for enhancing the synthesis of malonyl-CoA | This work<br>This work |
| Po1f/AT4S$_{x5}$I-EcACCABCD | Po1f/AT4S$_{x5}$I containing pYLXP'2-EcACCABCD, used for enhancing the synthesis of malonyl-CoA | This work |
| Po1f/AT4S$_{x5}$I-EcACCABCD-EcBirA | Po1f/AT4S$_{x5}$I containing pYLXP'2-EcACCABCD-EcBirA, used for enhancing the synthesis of malonyl-CoA | This work |
| Po1f/AT4S$_{x5}$I-YlACC1 | Po1f/AT4S$_{x5}$I containing pYLXP'2-YlACC1, used for enhancing the synthesis of malonyl-CoA | This work |

| | | |
|---|---|---|
| Po1f/AT4S$_{x5}$I-YlACC1-YlBPL1 | Po1f/AT4S$_{x5}$I containing pYLXP'2-YlACC1-YlBPL1, used for enhancing the synthesis of malonyl-CoA | This work |
| Po1f/AT4S$_{x5}$I-EcACS-YlACC1 | Po1f/AT4S$_{x5}$I containing pYLXP'2-EcACS-YlACC1, used for enhancing the synthesis of malonyl-CoA | This work |
| Po1f/AT4S$_{x5}$I-ScACS2-YlACC1 | Po1f/AT4S$_{x5}$I containing pYLXP'2-ScACS2-YlACC1, used for enhancing the synthesis of malonyl-CoA | This work |
| Po1f/AT4S$_{x5}$I-YlACS2-YlACC1 | Po1f/AT4S$_{x5}$I containing pYLXP'2-YlACS2-YlACC1, used for enhancing the synthesis of malonyl-CoA | This work |
| Po1f/AT4S$_{x5}$IHR$_{x2}$H-YlACS2-YlACC1 | Po1f/AT4S$_{x5}$I containing pYLXP'2-GhF3'H-CrCPR$_{x2}$-SlF3H-YlACS2-YlACC1, used for introducing Module II to Po1f/AT4S$_{x5}$I-YlACS2-YlACC1 | This work |
| NarPro/ASC | Po1f containing integrative naringenin pathway and over-expressing *YlARO1*, *YlACS2*, and *YlACC1* | (*12*) |
| ErioPro | Po1f containing integrative eriodictyol pathway and over-expressing *YlARO1*, *YlACS2*, and *YlACC1* | (*12*) |
| TaxiPro | Po1f containing integrative taxifolin pathway and over-expressing *YlARO1*, *YlACS2*, and *YlACC1* | (*12*) |

**Table S3. Primers used in this paper.**

| Primer | Sequence (5'-3') |
|---|---|
| RtTAL F | GACCAGCACTTTTTGCAGTACTAACCGCAGGCGCCTCGCCCGACTTCGCAAAGCC |
| RtTAL R | GGGACAGGCCATGGAACTAGTCGTTATGCCAGCATCTTCAGCAGAACATTG |
| Pc4CL F | GACCAGCACTTTTTGCAGTACTAACCGCAGGGTGACTGCGTTGCCCCGAAAG |
| Pc4CL R | GGGACAGGCCATGGAACTAGTCGTTACTTCGGCAGGTCGCCGCTCGCA |
| PhCHS F | GACCAGCACTTTTTGCAGTACTAACCGCAGGTTACGGTGGAAGAATACCGC |
| PhCHS R | GGGACAGGCCATGGAACTAGTCGTTAGGTAGCCACACTATGCAGAACC |
| MsCHI F | GACCAGCACTTTTTGCAGTACTAACCGCAGGCAGCAAGCATTACGGCAATCAC |
| MsCHI R | GGGACAGGCCATGGAACTAGTCGTCAGTTACCGATTTTAAAGGCACC |
| EcACS F | ACCAGCACTTTTTGCAGTACTAACCGCAGAGCCAAATTCACAAACACAC |
| EcACS R | CATAGCACGCGTGTAGATACTTACGATGGCATCGCGATAGCC |
| CgACC F | ACCAGCACTTTTTGCAGTACTAACCGCAGGTGTCAGTCGAGACTAGGAAG |
| CgACC R | CATAGCACGCGTGTAGATACTTACAGTGGCATGTTGCCGTGCTTGC |
| ARO1_Up F | GACCAGCACTTTTTGCAGTACTAACCGCAGTTTGCCGAGGGTCAGATCCAAAAGG |
| ARO1_Up R | TAGAATGCCGCAGTCTTAATGACCTCCGCCATACC |
| ARO1_Down F | GCGGAGGTCATTAAGACTGCGGCATTCTACGACGC |
| ARO1_Down R | GGGACAGGCCATGGAACTAGTCGTTAGTTACCAAGAACAGCCTTC |
| ScCPR1 F | CCAGCACTTTTTGCAGTACTAACCGCAGCCGTTTGGAATAGACAACACC |
| ScCPR1 R | CATAGCACGCGTGTAGATACTTACCAGACATCTTCTTGGTATC |
| YlCPR F | ACCAGCACTTTTTGCAGTACTAACCGCAGGCTCTACTCGACTCTCTCGAC |
| YlCPR R | CATAGCACGCGTGTAGATACCTACCACACATCTTCCTGGTAGAC |
| YlACS2 F | GACCAGCACTTTTTGCAGTACTAACCGCAGTCTGAAGACCACCCAGCCATC |
| YlACS2 R | TGCATAGCACGCGTGTAGATACTTACTTTTTCAACGAGTGAAC |
| ScACS2 F | GACCAGCACTTTTTGCAGTACTAACCGCAGACAATCAAGGAACATAAAGTAG |
| ScACS2 R | AGGCCATGGAACTAGTCGGTACCTTATTTCTTTTTTTGAGAG |

**Table S4. Plasmids used in this paper.**

| Plasmid | Annotation |
|---|---|
| pYLXP’ | YaliBrick plasmid, used for pathway assemble |
| pYLXP’2 | YaliBrick plasmid, used for pathway assemble |
| pYLXP’-Pc4CL-PhCHS-MsCHI | For the construction and optimization of Module I |
| pYLXP’-Pc4CL-PhCHS-PhCHI | For the construction and optimization of Module I |
| pYLXP’-Pc4CL-HvCHS2-MsCHI | For the construction and optimization of Module I |
| pYLXP’-Pc4CL-HvCHS2-PhCHI | For the construction and optimization of Module I |
| pYLXP’-Sl4CL-PhCHS-MsCHI | For the construction and optimization of Module I |
| pYLXP’-Sl4CL-PhCHS-PhCHI | For the construction and optimization of Module I |
| pYLXP’-Sl4CL-HvCHS2-MsCHI | For the construction and optimization of Module I |
| pYLXP’-Sl4CL-HvCHS2-PhCHI | For the construction and optimization of Module I |
| pYLXP’2-GhF3’H-CrCPR | For the construction and optimization of Module II |
| pYLXP’2-GhF3’H-YlCPR | For the construction and optimization of Module II |
| pYLXP’2-GhF3’H-ScCPR | For the construction and optimization of Module II |
| pYLXP’2-GmF3’H-CrCPR | For the construction and optimization of Module II |
| pYLXP’2-GmF3’H-YlCPR | For the construction and optimization of Module II |
| pYLXP’2-GmF3’H-ScCPR | For the construction and optimization of Module II |
| pYLXP’-T4SI | Plasmid pYLXP’-RtTAL-Pc4CL-PhCHS-MsCHI, used for the construction and optimization of Module I |
| pYLXP’-T$_{x2}$4SI[(1)] | Plasmid pYLXP’-RtTAL$_{x2}$-Pc4CL-PhCHS-MsCHI, used for the optimization of Module I |
| pYLXP’-T4$_{x2}$SI | Plasmid pYLXP’-RtTAL-Pc4CL$_{x2}$-PhCHS-MsCHI, used for the optimization of Module I |
| pYLXP’-T4S$_{x2}$I | Plasmid pYLXP’-RtTAL-Pc4CL-PhCHS$_{x2}$-MsCHI, used for the optimization of Module I |
| pYLXP’-T4SI$_{x2}$ | Plasmid pYLXP’-RtTAL-Pc4CL-PhCHS-MsCHI$_{x2}$, used for the optimization of Module I |
| pYLXP’-T4S$_{x3}$I | Plasmid pYLXP’-RtTAL-Pc4CL-PhCHS$_{x3}$-MsCHI, used for the optimization of Module I |
| pYLXP’-T4S$_{x4}$I | Plasmid pYLXP’-RtTAL-Pc4CL-PhCHS$_{x4}$-MsCHI, used for the optimization of Module I |

| | |
|---|---|
| pYLXP'-T4S$_{x5}$I | Plasmid pYLXP'-RtTAL-Pc4CL-PhCHS$_{x5}$-MsCHI, used for the optimization of Module I |
| pYLXP'2-HRH | Plasmid pYLXP'2-GhF3'H-CrCPR-SlF3H, used for the construction and optimization of Module II |
| pYLXP'2-H$_{x2}$RH | Plasmid pYLXP'2-GhF3'H$_{x2}$-CrCPR-SlF3H, used for the optimization of Module II |
| pYLXP'2-HR$_{x2}$H | Plasmid pYLXP'2-GhF3'H-CrCPR$_{x2}$-SlF3H, used for the optimization of Module II |
| pYLXP'2-HRH$_{x2}$ | Plasmid pYLXP'2-GhF3'H-CrCPR-SlF3H$_{x2}$, used for the optimization of Module II |
| pYLXP'2-HR$_{x3}$H | Plasmid pYLXP'2-GhF3'H-CrCPR$_{x3}$-SlF3H, used for the optimization of Module II |
| pYLXP'2-HRx4H | Plasmid pYLXP'2-GhF3'H-CrCPRx4-SlF3H, used for the optimization of Module II |
| pYLXP'2-HR$_{x5}$H | Plasmid pYLXP'2-GhF3'H-CrCPR$_{x5}$-SlF3H, used for the optimization of Module II |
| pYLXP'-AT4S$_{x5}$I | Plasmid pYLXP'-YlARO1-RtTAL-Pc4CL-PhCHS$_{x5}$-MsCHI, used for the optimization of Module I |
| pYLXP'2-CgACC | For enhancing malonyl-CoA synthesis |
| pYLXP'2-EcACCABCD | For enhancing malonyl-CoA synthesis |
| pYLXP'2-EcACCABCD-EcBirA | For enhancing malonyl-CoA synthesis |
| pYLXP'2-YlACC1 | For enhancing malonyl-CoA synthesis |
| pYLXP'2-YlACC1-YlBPL1 | For enhancing malonyl-CoA synthesis |
| pYLXP'2-EcACS-YlACC1 | For enhancing malonyl-CoA synthesis |
| pYLXP'2-ScACS2-YlACC1 | For enhancing malonyl-CoA synthesis |
| pYLXP'2-YlACS2-YlACC1 | For enhancing malonyl-CoA synthesis |
| pYLXP'2-HR$_{x2}$H-YlACS2-YlACC1 | For enhancing malonyl-CoA synthesis |

(1) The subscripts "$_{x2}$", "$_{x3}$", "$_{x4}$", and "$_{x5}$" refer to gene copy number in the plasmids.

### Supplementary figures

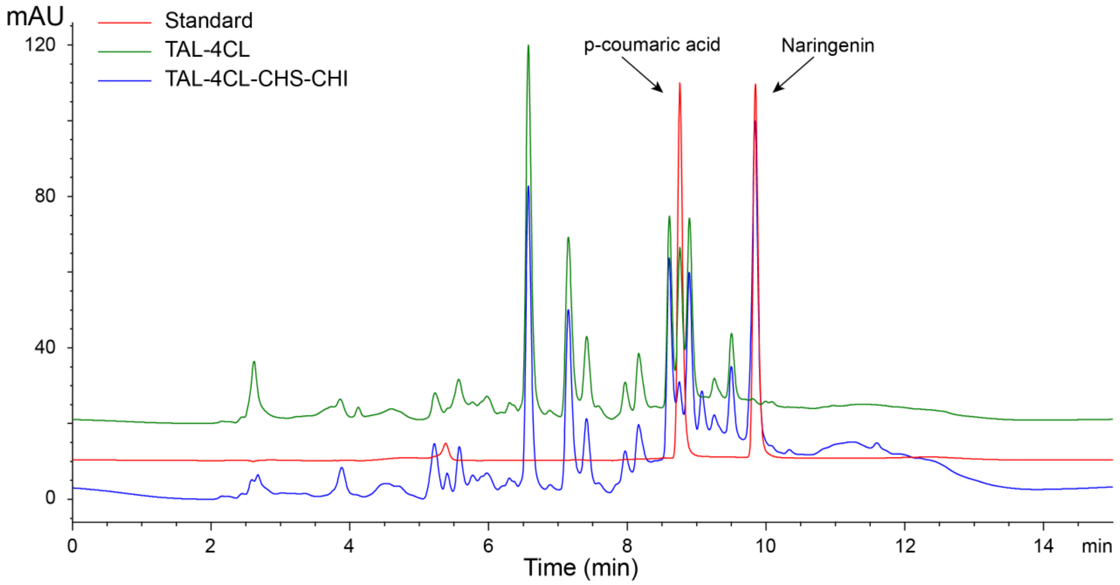


**Supplementary Figure S1.** HPLC profile of strains producing both naringenin and *p*-coumaric acid. Red lines, HPLC standards for *p*-coumaric acid and naringenin. Green line, strains with the overexpression of TAL and 4CL, only produced *p*-coumaric acid. Blue line, strains with the overexpression of TAL-4CL-CHS-CHI, produced both *p*-coumaric acid and naringenin.

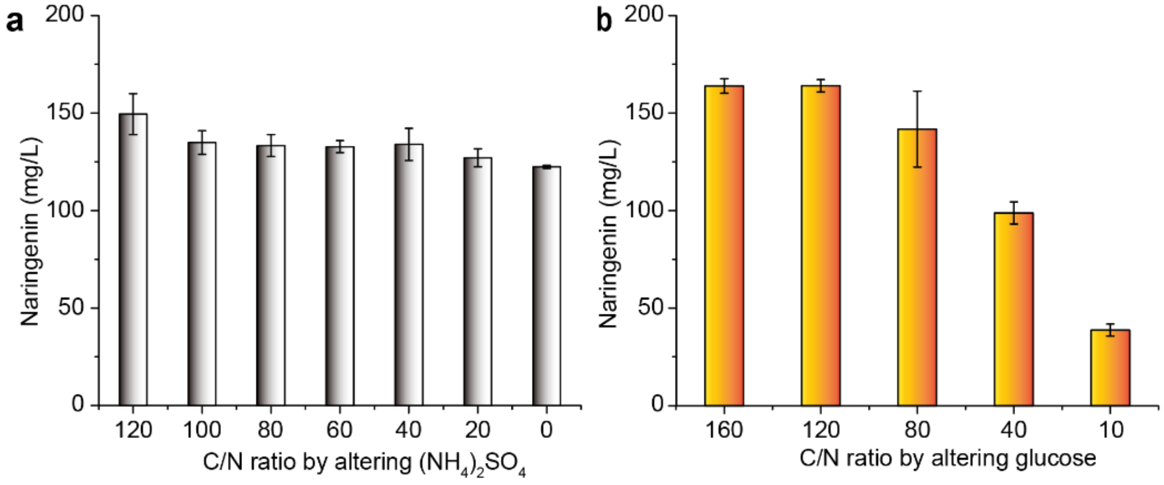


**Supplementary Figure S2.** C/N ratio optimization using different patterns. **a.** C/N ratio optimization by fixing glucose content and altering $(NH_4)_2SO_4$ content. **b.** C/N ratio optimization by fixing $(NH_4)_2SO_4$ content and altering glucose content. Both optimizations were carried out in leucine and uracil drop-out complete synthetic media (CSM-Leu-Ura).

Because substantial carbon flux was directed to lipid biosynthesis in the oleaginous yeast *Yarrowia lipolytica*, it is believed that sufficient oleic acid supplement will inhibit lipid biosynthesis and thus redirect carbon flux to flavonoid biosynthesis. Acetate can be used as the precursor for cytosolic acetyl-CoA and malonyl-CoA (*13*). Cerulenin is an inhibitor of lipid acid biosynthesis pathway (*14*). In the subsequent optimizations, the effects of oleic acid, sodium acetate, and cerulenin were investigated. The results showed that adding cerulenin improved naringenin titer by 53.2%, reaching 109.0 mg/L, while neither oleic acid nor sodium acetate had obvious effect on naringenin titer (Supplementary Fig. S3).

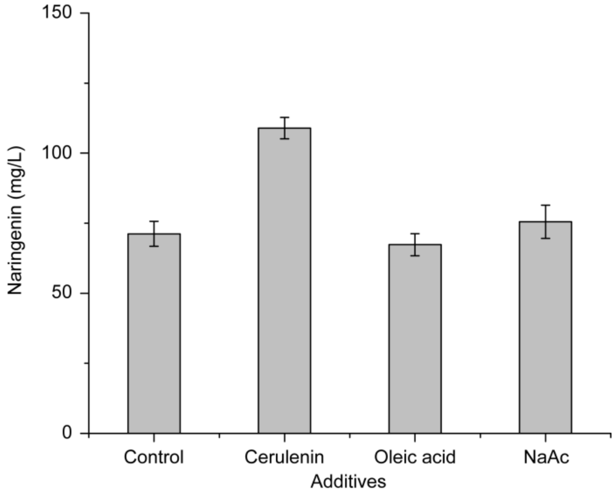


**Supplementary Figure S3.** Effects of cerulenin, oleic acid, and sodium acetate on naringenin titer. The final concentrations of cerulenin, oleic acid, and Na-Ac were 1 mg/L, 5 g/L, and 2 g/L respectively. Oleic acid and NaAc were added at the start of the fermentation, while cerulenin was added at 48 h.

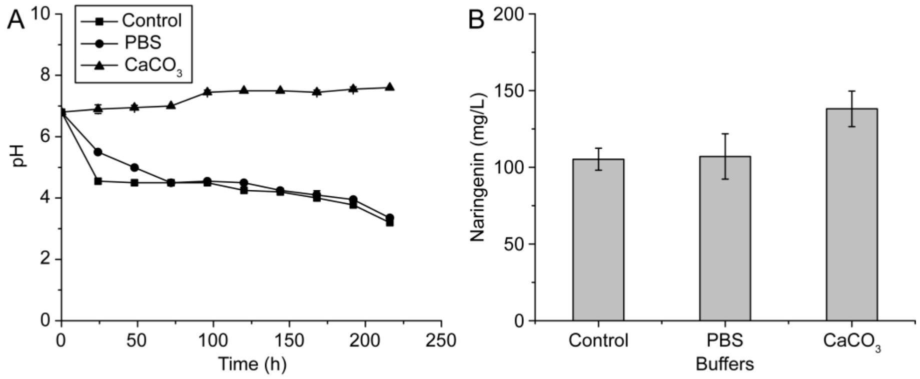


**Supplementary Figure S4.** Effects of PBS and $CaCO_3$ on the pH and naringenin titer. **A.** Time course of the pH during the fermentation process. Twenty mM PBS or 40 g/L $CaCO_3$ was used to buffer the acidity. **B.** Effects of PBS and $CaCO_3$ on naringenin titer. The naringenin titer was measured at 144 h.

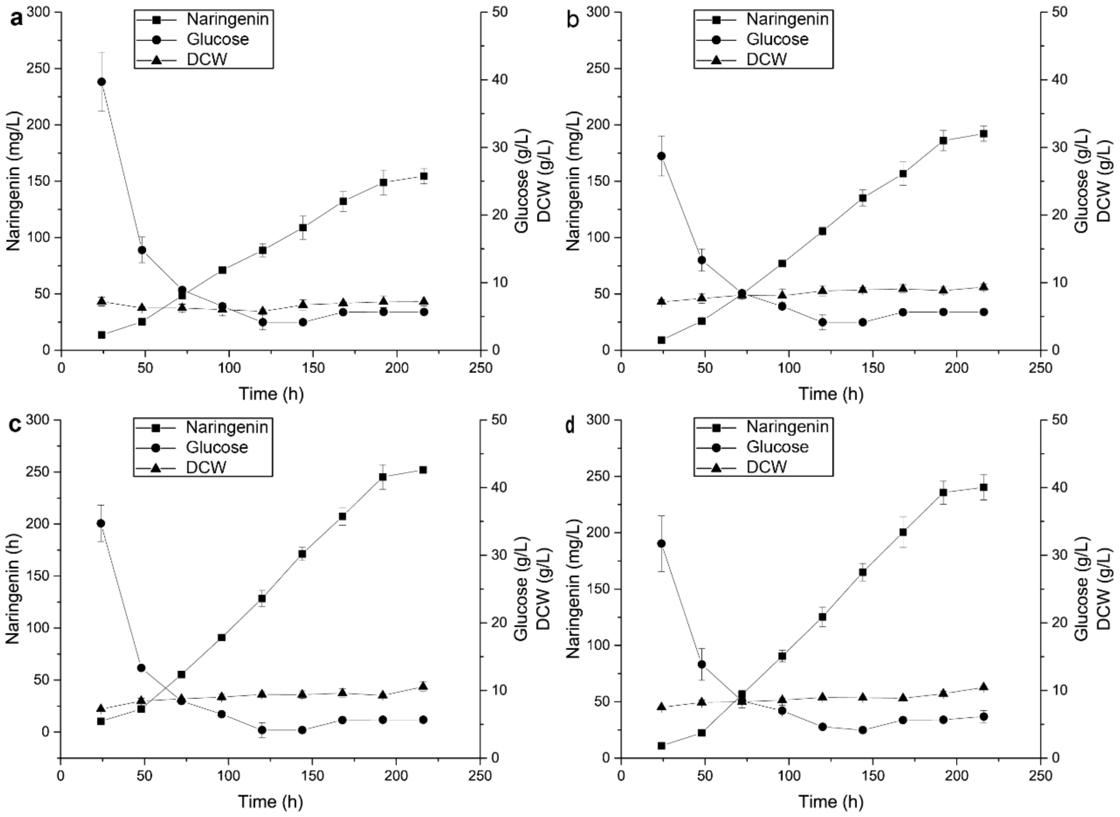


**Supplementary Figure S5.** Time course of fed-batch fermentation in YPD media. **a.** Control. **b.** Buffered with 40 g/L $CaCO_3$. **c.** Buffered with 40 g/L $CaCO_3$. A final concentration of 1 mg/L cerulenin was added at 48 h. **d.** Buffered with 40 g/L $CaCO_3$. A final concentration of 1 mg/L cerulenin was added at 48 h, and a final concentration of 5 mM NaAc was added at 24 h intervals. For all experiments, the starting glucose concentration was 40 g/L, and 10 g/L glucose was added at 24h and 48 h.
